# Unveiling Urban Complexity: Research note on integrating OpenStreetMap to enhance representation of fine-scale landscape heterogeneity

**DOI:** 10.1101/2023.10.31.564785

**Authors:** Tiziana A. Gelmi-Candusso, Peter Rodriguez, Marie-Josée Fortin

**Author notes:** **Corresponding Author** Tiziana A. Gelmi Candusso, PhD.

## Abstract

Landscape heterogeneity has an impact on wildlife behavior, their interactions, and their persistence. Urban landscapes are among the world’s most heterogeneous landscapes, yet current global landcover maps classify developed land in a single landcover type. This limits the spatial scale at which urban ecologists can approach research questions. OpenStreetMap (OSM), an open-source mapping platform, can be leveraged to enhance the representation of landscape heterogeneity in developed areas. For this, we extracted OSM features with attributes representing infrastructure, land use and green cover, integrating these into a continental landcover map through a globally applicable computational framework. We validated our OSM-enhanced landcover layer against existing remote sensing, aerial photography, and local governmental maps for 33 cities in North America. Our framework’s output provides an 89% accurate representation of landscape heterogeneity. We discuss caveats, potential improvements, and ecological applications. Our OSM-based landcover enhancement framework will facilitate the use of open-source landscape information for improved ecological modeling and urban planning.

## Introduction

Urban landscapes are composed of a heterogeneous matrix of diversely built environments, vegetated areas with different management strategies, and linear features with varying types of usage and degrees of traffic. This landscape heterogeneity can strongly influence the spatial ecology of local wildlife, their habitat use, and behavior (App et al., 2022; Hansen et al., 2020; Wurth et al., 2020). However, the urban matrix is mostly presented as a homogenous “developed”/”built” class in global land use/landcover (LULC) maps (ESA, 2017; Karra et al., 2021). Such an approach falls short in capturing the landscape fragmentation that shapes ecological processes, resulting in a diminished relevance and applicability of these landcover maps within the context of urban ecology.

To overcome this issue, most urban studies analyzing habitat use and wildlife behavior rely on LULC maps provided by local governmental agencies. These include finer landscape information such as land use classes, which provides a representation of human activity across the landscape (Randa & Yunger, 2006; Teitelbaum et al., 2020; Thompson et al., 2021), building footprints, and centerline layers representing road networks. Yet, these local LULC maps are highly sensitive to sampling bias and might not be equivalent between cities, limiting multi-city comparisons (Thomlinson et al., 1999). Furthermore, the availability of these maps may be limited in countries with lower mapping resources, excluding a priori the inclusion of certain study areas in global analysis. While remote sensing for urban landcover classification is making remarkable advances at the global scale (Kuras et al., 2021), and there have been advances toward a finer-scale representation of urban heterogeneity using aerial photography and fine resolution satellite products (Cadenasso et al., 2007; Pesaresi & Politis, 2023a). The suggested methodologies have low accessibility given computational demands and high costs or have limited representation of human activities across the landscape – i.e. land use – a key aspect in urban ecology (Gallo et al., 2022). Therefore, an open-source and reproducible approach to generate a fine classification of the urban landscape, which includes both the natural and human components, is imperative to increase accessibility and comparability across cities for future research.

OpenStreetMap (OSM), an open-source mapping platform, provides fine vector-based landscape information characterizing landscape heterogeneity (OpenStreetMap contributors, 2017). These data are generated by community members and supplemented with data from governmental and non-governmental agencies around the globe. OSM has been previously used in ecological research for extracting settlements and roads (Bide et al., 2023; Gizatullin & Alekseenko, 2022), extracting elevation (Shaykevich et al., 2022), distance to water sources (de la Torre et al., 2021), and to study the fractal dimension of cities (Malishevsky, 2022). However, a standardized framework to extract all attributes characterizing urban features and develop accurate landcover data layers for ecological analysis and urban planning, has yet to be proposed.

Hereby, we propose a globally applicable computational framework to generate a fine representation of urban landscapes by integrating OSM features into a global landcover map. We validate our OSM-based enhancement framework by quantifying its completeness and accuracy across 33 cities. We use building features as a case study to compare our framework against available remote sensing layers and local building footprints derived from aerial photography and LiDAR. The output of this computational framework will increase accessibility to the information needed to integrate the fine landscape heterogeneity intrinsic of urban areas into ecological studies and urban planning.

## Methods

### OSM attribute extraction

The OSM database consists of a series of features represented in space as points (nodes), lines (ways), and polygons (relations). Each feature is associated with a series of descriptive attributes (known as tags) which consist of key-value pairs (e.g., land use: residential). We systematically queried the OSM database, using the OSM dictionary and visualization tool, identified and extracted all features with attributes describing components within the urban landscape and its surroundings. (OpenStreetMap contributors, 2023) (Table S1).

We categorized the extracted OSM features into 27 landcover classes following their descriptive attributes, distributing features across three categories (Table S1, Table S2): (i) human land use, encapsulating human activities classified by type (e.g., residential, commercial, industrial, institutional); (ii) green cover, classified based on vegetation density and management approach, we included here also water cover and barren soil; and (iii) infrastructure, encompassing physical structures like buildings, fences, and linear features such as railways and roads classified by road type and hence their hypothetical traffic load. While features represented as polygons (relations) could be extracted as is, linear features (ways), such as roads, necessitated conversion into polygons using buffers of varying sizes following feature type and observed mean lane number (Table S3).

### Overlaying OSM attributes

Following the extraction and classification of the OSM vector-based features, we converted the extracted polygons into 30m resolution raster-based features using the rasterize() function from the *terra* package in R (R Core Team, 2013). We rasterized to reduce computational resources required, given the large extent of our study areas, and to facilitate the integration of the OSM features into the global landcover map used hereby, however for smaller areas, and other global landcover maps, the resolution can be modified within the framework. Subsequently, to consolidate the 27 classes of OSM features into a single map layer, we superimposed the raster-based features according to a priority structure (Table S1), overlaying the green cover layers over the land use layers, and on top of these the infrastructure layers. Within the infrastructure layers, we overlaid buffered roads over buildings, layered by road type and traffic load optimizing for overpasses and underpasses, and finally above all features, we overlaid hiking paths, railways, and fences.

### Integration of OSM features into global landcover map

To integrate the OSM features into the global landcover layer, we reclassified the latter to our 27-class landcover classification approach (Table S4), introducing a new “developed land” class (the 28th class) to preserve continental-scale undefined classification of urban areas and prevent misrepresenting human activity (Figure 1, Table S1). Any raster cells in the OSM-based map layer that was devoid of information were characterized following their value on the reclassified continental-scale landcover value. Hereby we opted for the CEC landcover data specific for North America (Commission for Environmental Cooperation (CEC), 2023), given our focus on North American cities. For intercontinental global applications, Copernicus (Buchhorn et al., 2020) or ESA WorldCover (Zanaga et al., 2021) raster layers maps are recommended.

**Figure 1.**
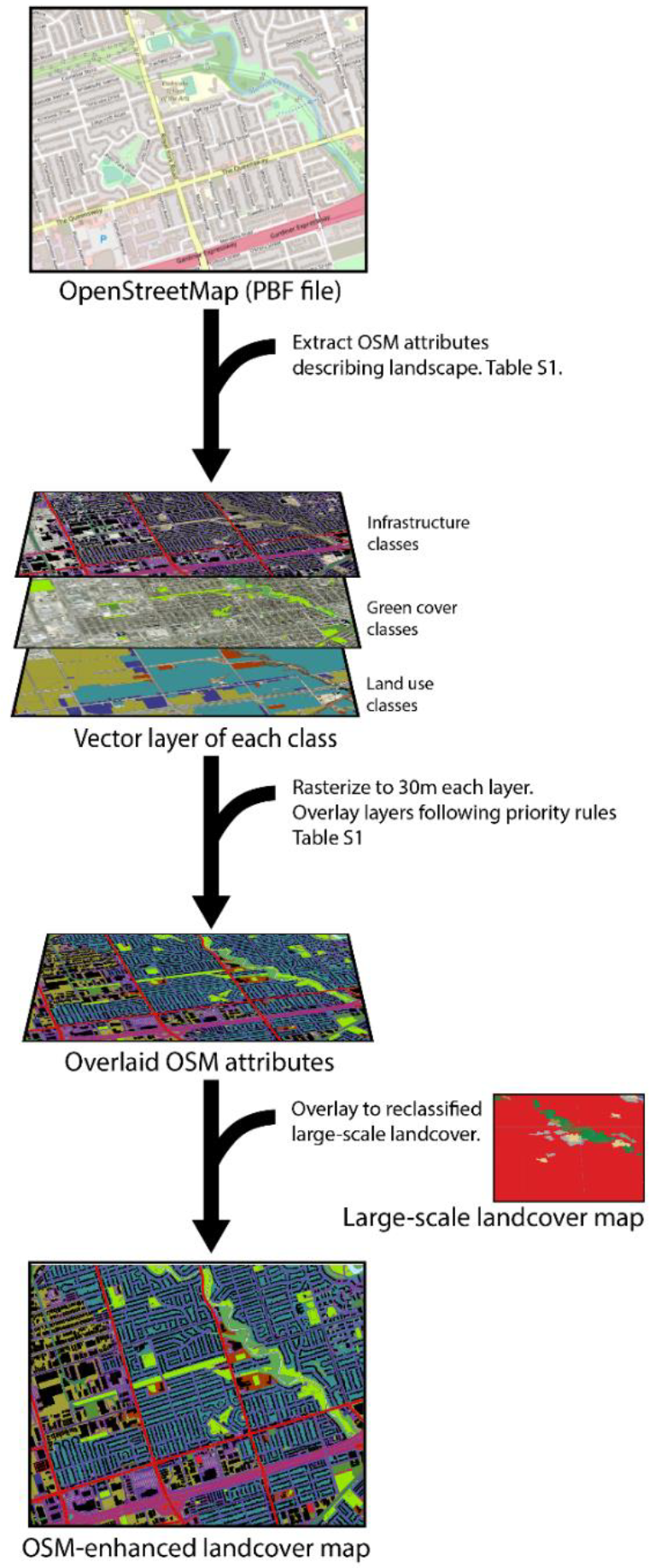
Figure 1 Workflow for enhancing the resolution of developed areas in global landcover maps using OpenStreetMap (OSM) attributes.

### Validation

We validated our framework’s results by assessing two measures: completeness and accuracy for 33 study areas encompassing North American cities. The study areas were defined by an envelope centered around each city’s administrative boundary that expanded up to an arbitrarily selected buffer size (Table S5). We assessed the completeness of the OSM attributes included in our framework by investigating the number of OSM cells devoid of data, estimating the absolute change in area, across landcover classes, between a map layer including only the extracted OSM features and the global map layer. We assessed the accuracy of the framework’s final output, by cross-validating 2123 sampling points stratified by landcover classes, against street view imagery and aerial photography (©2023 Google). As accuracy measures, we estimated the Kappa statistic (i.e. agreement rate between two datasets given the agreement expected by random coincidence (Monserud & Leemans, 1992)), and the precision of each class (i.e. the rate at which each class identified correctly the landscape) using the confusionMatrix() function from the *caret* package.

### Case study: Building footprint

To investigate in detail how extracting information from OSM compares to using information provided by remote sensing products against local mapping products, we further analyzed the buildings class, an intrinsic infrastructure of urban areas and potentially underrepresented class across layers. The remote sensing products we compared against were open-access global layers sourced from the Global Human Settlement project (GHL) and included (1)a layer including building information in terms of building density (GHL-S, 10m resolution), and (2) a layer including the building footprint (GHL-MSZ, 10m resolution) (Pesaresi & Politis, 2023b). As a reference, we used local vector-based building footprint layers provided by respective cities which were either generated computationally from fine resolution landcover layers, from aerial photography or LiDAR (Light detection and ranging), rasterized to 30m (Table S6). We quantified the differences in proportion of total building area estimated by each mapping system for 20 of our 33 study areas, where a local building footprint layer was available (Table S6).

## Results

Our framework augmented the “developed/built” areas of the global landcover map, by introducing the characteristic urban landscape heterogeneity stemming from varying human activities, building densities, and transportation networks within urban areas (Figure 2.1).

**Figure 2.**
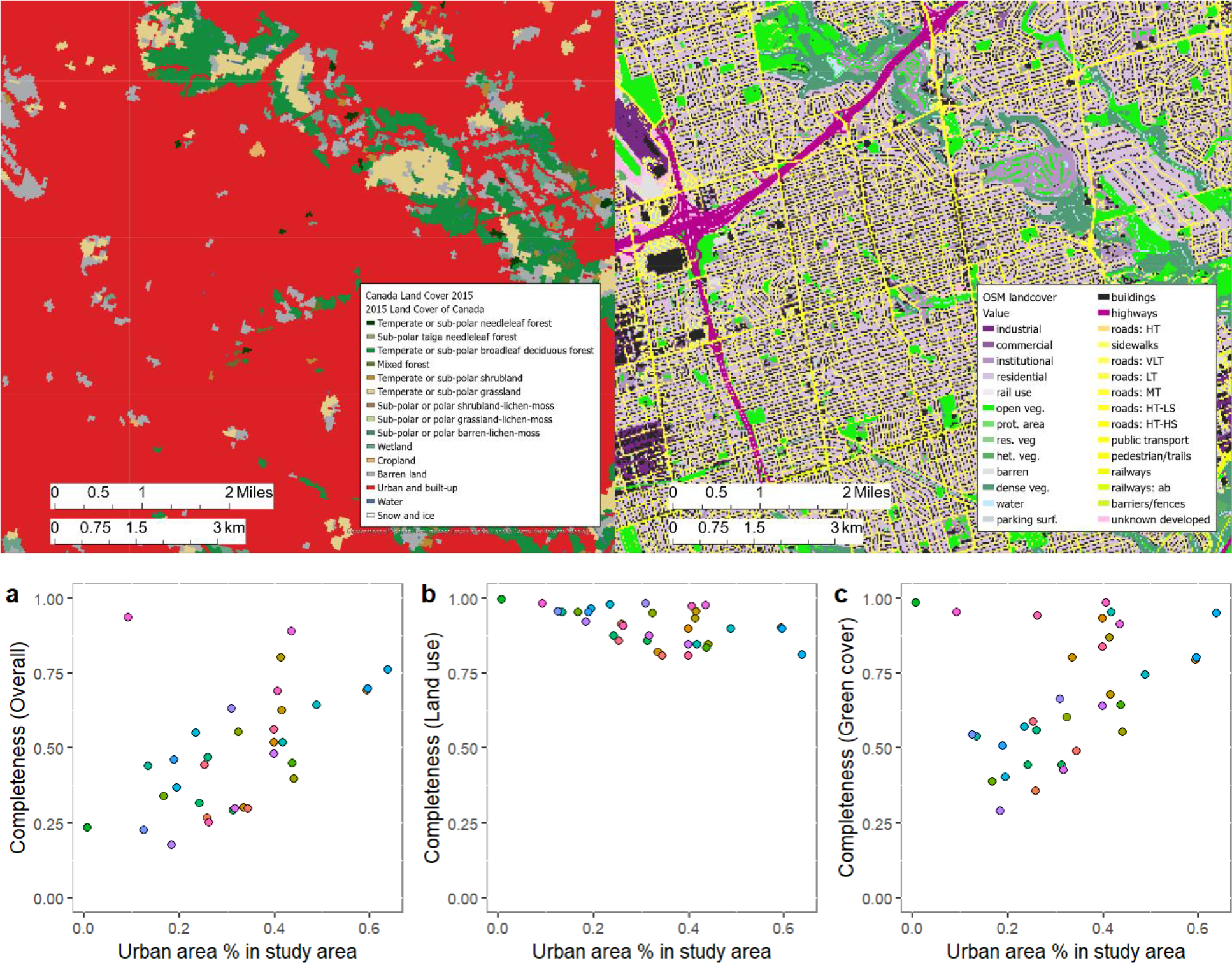
(i) Comparison global landcover vs OSM-augmented landcover. (ii) Completeness of the OSM-derived landcover map with increasing proportion of the surface area occupied by developed landcover (i.e. land use and infrastructure). a. Total completeness, includes all landcover types. b. Completeness of land use landcover types including human activities (e.g. industrial, commercial, residential) and c. Completeness of green cover types excluding water.

Across study areas, the completeness of the OSM database consistently increased as the urban extent within the study area increased (Figure 2.2). When distinguishing between landcover class groups, this effect was seen mainly for the green cover classes (50-100%, Figure 2.2b), especially forested areas (Figure S1). Instead, the completeness of land use classes was high (80-100%) throughout the cities, regardless of proportion of urban area included (Figure 2.2c). As a result, study areas with larger envelope buffer size, for example, San Diego County and Maryland, had the lowest completeness values. The completeness of water was low only for coastal cities, as these included a large body of open water in the study area, not generally represented in the OSM database (Figure S1).

The percentage of areas occupied by classes with undefined sub-tipology stemming from undefined attributes in the OSM database was low. Only a median of 1.6% of the total road-classified study area lacked road type information (Figure S2a) and a median of 0.2% of the total surface area occupied by vegetation landcover classes lacked information on vegetation type. (Figure S2b).

In terms of accuracy, our framework’s final output had a Kappa statistic of 0.89. The classes with lower precision rates, i.e. how often the description of the cell was correct, were water (class 12, *Precision* = 0.78), linear features in construction or abandoned (class 26, *Precision* = 0.56) and the grouped developed/built class directly obtained from the continental scale landcover map when OSM cells were void (class 28, *Precision* = 0.35) (Table S7). The false positives in these classes included dried rivers, overgrown vegetation, and urban green cover types, respectively (Figure S3a). When considering only classes with unclassified sub-tipology, we found unclassified roads were predominantly residential (low traffic) or service roads (very low traffic), protected areas were predominantly dense, heterogeneous or open green cover, and unclassified developed areas were predominantly residential or some type of green cover (Figure S3b).

When comparing building information provided by remote sensing and OSM against local building footprints generated using LiDAR or orthophotos, our framework integrating OSM estimated a total building-occupied area closer to the local building footprints provided by governmental agencies. Remote-sensing products instead overestimated the building-occupied area across cities, when compared to local building footprints (Figure 3).

**Figure 3.**
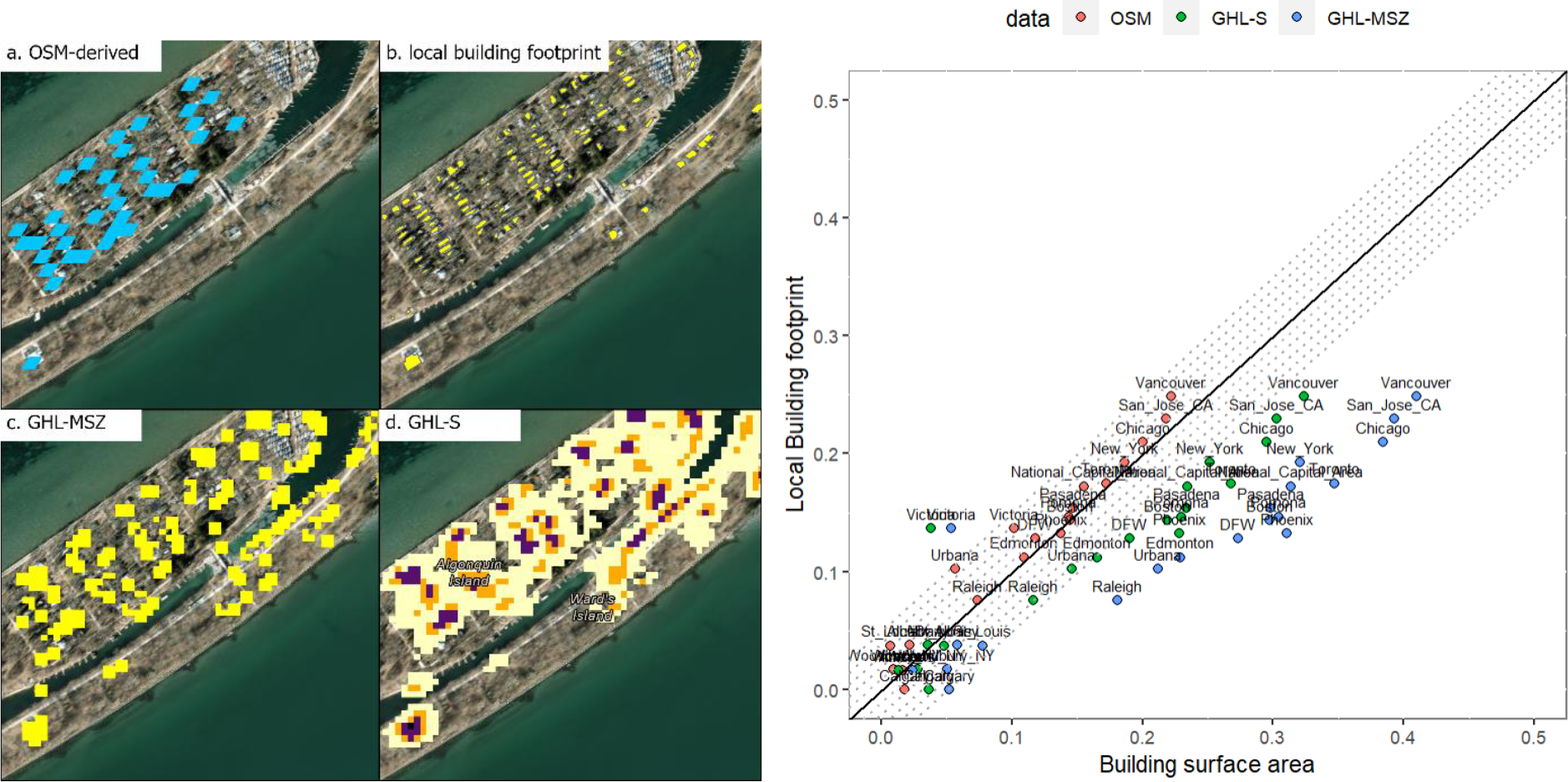
(i) Example of buildings represented by the (a) OSM building layer, (b) local building footprint, and remote sensing products, i.e. (c) GHL-MSZ and (d) GHL-S. (ii) Comparison between the total proportion of building surface estimated by the OSM-augmented landcover (OSM) and the Global human settlement remote-sensing building products (GHL-S, GHL-MSZ).

## Discussion

The integration of the hereby extracted OSM attributes into a global landcover layer accurately increased the representation of urban landscape heterogeneity. These OSM attributes added transportation networks, building footprints, vegetation cover intensity and human land use information to the otherwise homogenous developed class in the global landcover map. In terms of vegetation cover, completeness of these classes in the OSM database diminished in the urban periphery, however, these are generally highly accurately represented in global landcover map used hereby (77.1-86.9%), as well other landcover maps potentially used instead, such as the ESA WorldCover (76.1%) or Copernicus global landcover map (80.6%). Given our findings, the importance of using background global landcover maps increases with the decreasing proportion of non-urban surface included in the study area. Furthermore, while we expected buildings to be the most underrepresented class in the OSM database, given the high number of these and the effort needed to add each individual polygon into the OSM database, OSM outperformed remote sensing products when compared to local building footprints using Orthophotos and LiDAR.

We found some limitations that should be considered when applying OSM to ecological studies. These included (1) linear features with incomplete and inconsistent information on lane numbers, crucial for buffering roads, (2) lack of information on vegetation type on private green spaces, such as yards, and (3) sometimes sparse information on land use areas, up to 20% of total area being unclassified especially for more rural cities.

The OSM-derived road network contained information on the number of lanes which we used as a reference to buffer these originally line vector-based features. However, lane numbers were not always included, and numbers and units were used inconsistently, which led us to ultimately use a buffer of 6m estimate a reference lane number for each road type, derived from observations in aerial photography and mean lane number in OSM. Being able to systematically and accurately buffer linear features to a representative diameter improves the applicability of OSM to ecological studies on animal movement and road mortality. In the absence of such information, road type, e.g. primary vs tertiary, which we use to approximate traffic load, can help overcome the lane number caveat, as our analysis shows road type information is rarely missing in the OSM database.

The OSM database does not include information on yard size area nor vegetation type within. Most private green areas are classified as residential. However, given the fine landscape representation of our framework, a logical assumption would be to consider areas not overlapped by buildings, parking lots, nor linear features, as residential green areas. These may then be overlaid to a vegetation index when applying the framework to study subjects strongly affected by yard vegetation complexity, e.g. hedgehogs (App et al., 2022), similar to open areas are overlaid to the NDVI in the GHL-c map (Pesaresi & Politis 2023a).

Certain areas were defined as protected areas in the OSM database, mainly natural parks, sometimes without further information on vegetation types. Fortunately, the total proportion of green areas occupied by these undefined green areas was generally below 1% across study areas. However, it is important to consider that these areas might affect ecological studies requiring information on how much shelter and food resource a green area provides. A potential solution to overcome such limitation is to replace the protected area class with the continental scale landcover reclassified information on vegetation type or a form of remote-sensing derived vegetation index.

In certain study areas, land use information was sparse, reaching up to 20% of total study area where information on human activity was absent (Figure 2.2b). This might be problematic for ecological studies focusing on animal species with a high degree of human avoidance which might differentially favor different land use types, e.g. coyotes (Thompson et al., 2021). To overcome this, efforts should be directed towards completing land use information in the OSM database. We suggest local verification of the percentage of missed land use area, by quantifying the unclassified developed area (class 28), when applying the framework to areas outside those tested here. Further improvement of our framework may include the inclusion of other potential ways of defining land use in the OSM such as using the boundaries layer instead of polygons, a distinct layer in the OSM database structure (e.g. Boston).

Compared to other map layers aiming at enhancing the representation of landscape heterogeneity in urban areas (Cadenasso et al., 2007; Pesaresi & Politis, 2023a), OSM increases the accessibility of fine resolution features and provides additional details not included in previous methods. For example, in comparison to the GHL MSZ layer (Pesaresi & Politis, 2023a), included in our case study, OSM integrates to the building footprint the transportation network, human activity outside of buildings, and represents all larger urban green areas, e.g. cemeteries, larger parks and urban forests (Figure S4). In comparison to the Hercules framework (Cadenasso et al., 2007), instead of relying on aerial photography and LiDAR, our framework relies on open access databases readily available across the globe, increasing the accessibility and applicability of fine resolution landscape information.

The integration of fine resolution landscape heterogeneity into landcover maps will provide key information to ecological studies, in particular those using agent-based or connectivity models involving fine resolution movement decisions within highly fragmented landscapes. Agent-based models integrating landscape have proven useful for understanding navigation patterns of Caribou in highly industrialized rural areas and Jaguars (*Panthera orca*) in protected areas in Belize, and for understanding landscape-dependent disease transmission in urban areas (Semeniuk et al., 2012; Watkins et al., 2015, Dimitrov et al., unpublished data). Connectivity models integrating fine resolution landcover maps, have shown yard vegetation complexity are important for hedgehogs (Braaker et al., 2014) and roads are important for red fox movement across the city and its resulting genetic connectivity (Kimmig et al., 2020). Furthermore, fine resolution landcover maps have been used to understand temporal changes in habitat use (Thompson et al., 2021), better explain biodiversity of invertebrates in cities (Savage et al., 2015), and improve the estimation of urban ecosystem services, such as total carbon storage (Grafius et al., 2016). Landscape heterogeneity was key within these studies, for which they relied on local landcover layers. The integration OSM will allow to overcome geographic boundaries imposed by mapping product availability, opening the opportunity for similar studies on a global scale, allowing for multi-city comparisons across countries and the inclusion of study areas where governmental mapping resources are scarce.

## Conclusion

The framework proposed in this study is able to integrate OSM features into widely used global landcover layers, providing a promising approach for revealing the fine-scale landscape heterogeneity of urban areas. The output is an improvement from other frameworks aiming at representing urban areas, primarily in terms of accuracy, accessibility, and geographic scale of applicability. We unravel limitations, potential solutions, and considerations for implementing the proposed framework to ecological studies. Studies benefiting from integrating a fine representation of landscape heterogeneity include modeling approaches investigating animal movement and behavior, epidemiological analyses, and landscape connectivity. For these, augmenting global landcover maps by integrating OSM features will enable worldwide multi-city comparisons and allow for the inclusion of commonly underrepresented areas, as it reduces the reliance on maps generated by local governments.

## Acknowledgements

We acknowledge and appreciate the contribution in the verification of the OSM-augmented landcover layer by the respective local partners at the Urban Wildlife Information Network (UWIN), Joaquin Rodriguez for the blind cross-validation of random samples using aerial photography and street view imagery, and the contribution validating OSM attributes by Juan De Los Rios and Janice Hau. We acknowledge and are grateful for the funding received from the School of Cities at the University of Toronto, the German Research Foundation (DFG), the Natural Sciences and Engineering Research Council of Canada (NSERC) and the Canada Research Chairs (CRC).

