## Supplementary material for "Unveiling Urban Complexity: Research note on integrating OpenStreetMap to enhance representation of fine-scale landscape heterogeneity": Table S1

Ecology and Evolutionary Biology department. University of Toronto. 25 Willcocks St. M5S3B2. Toronto (ON), Canada

**Supplementary material**

**Supplementary Tables**

Table S1. Landcover classes included in the OSM-derived urban fine-scale landcover map, including priority values and tags extracted from OSM for each category, land use, green cover and infrastructure.

| | | | Attributes (tags) | |
|---|---|---|---|---|
| | **Priority** | **Feature** | **Key** | **Value** |
| Land use | 1 | Industrial | land use | industrial, fairground |
| | | | industrial | factory |
| | 2 | Commercial | land use | commercial, retail |
| | 3 | Institutional | land use | institutional, education, religious, military |
| | | | amenity | school, hospital, university, fast_food, clinic, theatre, conference_center, place_of_worship, police |
| | | | leisure | golf_course |
| | | | healthcare | clinic, hospital |
| | 4 | Residential | land use | residential |
| | 5 | Land use: rail | land use | railway |
| | 28 | Developed: unclassified | NA | NA |
| Green cover | 6 | Open green area | land use | park,grass, cemetery, greenfield, recreation_ground, winter_sports |
| | | | golf | not rough |
| | | | amenity | park |
| | | | leisure | park, stadium, playground, pitch, sports_centre, stadium, pitch, picnic_table, pitch, dog_park, playground |

| | | | | |
|---|---|---|---|---|
| | | | sport | soccer |
| | | | power | substation |
| | | | surface | grass |
| | 7 | Protected area | leisure | nature_reserve |
| | | | boundary | protected_area, national_park |
| | | | protected_ area | nature |
| | | | land use | nature_reserve, natural_reserve, landscape_reserve |
| | 8 | Resourceful/a gricultural area | land use | orchard,farmland, landfill,vineyard, farmyard, allotments, allotment, farmland |
| | | | leisure | garden |
| | | | allotments | all |
| | 9 | Heterogenous green area | natural | garden, scrub, shrubbery, tundra, cliff, shrub, wetland, grassland, fell, heath,moor |
| | | | land use | plant_nursery, meadow, flowerbed, wetland |
| | | | meadow | all |
| | | | golf | rough |
| | | | grassland | pairie |
| | 10 | Barren soil | natural | mud, dune, sand, scree, sinkhole, beach |
| | | | land use | brownfield, construction |
| | | | golf | bunker |
| | 11 | Dense/foreste d green area | land use | forest |
| | | | natural | wood |
| | | | boundary | forest, forest_compartment |
| | 12 | Water | land use | basin |
| | | | natural | water, spring, waterway |
| | | | waterway | river, streat tidal_channel, canal, drain, ditch, yest |
| | | | water | all, not intermittent |
| | | | basin | detention |
| | | | intermitte nt | not yes |
| | | | seasonal | not yes |
| | | | tidal | not yes |
| Infrastructure | 13 | Parking surface | parking | surface |
| | | | aeroway | runway, apron |
| | 14 | Building | building | all, hospital, parking, industrial, school, commercial, terrace, detached, semideatched_house, house, retail, hotel, apartments, yes, airport, university |
| | | | parking | multi-storey |
| | | | aeroway | terminal |
| | 15 | Roads: very high traffic | highway | motorway,motorway_link, motorway_junction |

| | | | | |
|---|---|---|---|---|
| | 16 | Roads: sidewalk | footway | sidewalk |
| | 17 | Roads: unclassified | highway | not (footway,construction,escape,cycleway,steps,bridleway,construction,path,pedestrian,track,abandoned, turning_loop,living_street, bicycle road, cyclestreet, cycleway lane,cycleway tracks, bus and cyclists, service,services, busway, sidewalk, residential, rest_area, primary, motorway_junction, secondary, secondary_link, tertiary, tertiary_link, motorway,motorway_link,trunk_link, trunk, corridor,elevator,platform,platform,crossing,proposed, razed) |
| | 18 | Roads: very low traffic | highway | services,service,turning_loop,living_street |
| | 19 | Roads: low traffic | highway | residential, rest_area, busway |
| | 20 | Roads: medium traffic | highway | tertiary, tertiary_link |
| | 21 | Roads: high traffic, low speed | highway | primary, primary_link, secondary, secondary_link |
| | 22 | Roads: high traffic, high speed | highway | trunk, trunk_link |
| | 23 | Public transport: streetcars/trams | railway | tram |
| | 24 | Roads: pedestrian/trails | highway | footway,construction,escape, cycleway,steps,bridleway,path,pedestrian,track, abandoned,bicycle road, cyclestreet, cycleway lane, cycleway tracks, bus and cyclists |
| | | | footway | not sidewalk, not crossing |
| | 25 | Public transport: railway | railway | light_rail,narrow_gauge,rail,preserved,tram |
| | | | railway | not tram |
| | 26 | Public transport: disused railway/highway | railway | abandonded, construction, disused |
| | | | highway | construction |
| | 27 | Barriers | barrier | all |

Table S2. Description of each landcover class included in the OSM-augmented landcover map.

| | PRIORITY | FEATURE | DESCRIPTION |
|---|---|---|---|
| LAND USE | 1 | Industrial | Areas occupied by factories and other manufacturing companies |
| | 2 | Commercial | Areas occupied by commercial areas or retail areas where commercial activities happen |
| | 3 | Institutional | Areas occupied by universities or hospitals or other institutions |
| | 4 | Residential | Areas occupied by residential buildings mainly, where people live |
| | 5 | Land use: rail | landscape area exclusive for railway use |
| | 28 | Developed: unclassified | land use not available |
| GREEN COVER | 6 | Open green area | parks, landscaped areas, grass areas |
| | 7 | Protected area | protected areas with unclassified vegetation type |
| | 8 | Resourceful/agricultural area | green areas with fruits or vegetables |
| | 9 | Heterogenous green area | non managed green area |
| | 10 | Barren soil | areas without vegetation nor concrete. e.g. soil or sand |
| | 11 | Dense/forested green area | forest, natural areas |
| | 12 | Water | water bodies |
| INFRASTRUCTURE | 13 | Parking surface | parking lots, only those on the surface, does not include parking complexes nor underground |
| | 14 | Building | Buildings, houses, etc. |
| | 15 | Roads: very high traffic | very high traffic volume, high speed, e.g. highways |
| | 16 | Roads: sidewalk | sidewalks |
| | 17 | Roads: unclassified | unclassified roads |
| | 18 | Roads: very low traffic | very low traffic volume, e.g. laneways |
| | 19 | Roads: low traffic | low traffic volume, e.g. residential streets |
| | 20 | Roads: medium traffic | medium traffic volume |
| | 21 | Roads: high traffic, low speed | high traffic volume, low speed |
| | 22 | Roads: high traffic, high speed | high traffic volume, high speed |

| | | | |
|---|---|---|---|
| | 23 | Public transport: streetcars/trams | streetcars, trams, usually at the same level as roads, does not include underground structures. |
| | 24 | Roads: pedestrian/trails | linear features without vehicular traffic, mainly trails and walking paths |
| | 25 | Public transport: railway | railway tracks |
| | 26 | Public transport: disused railway/highway in construction | abandoned, unused trailway tracks where vegetation has overgrown or barren land where linear features are being built |
| | 27 | Barriers | any physical barrier to movement, fences, walls, etc. |

Table S3. Buffers used to convert OSM line features into polygons.

| **Layer name** | **Buffer (m)** |
|---|---|
| pedestrian, trails | 3 |
| sidewalks | 3 |
| roads: very low traffic | 6 |
| roads: low traffic | 12 |
| roads: medium traffic | 12 |
| roads: high traffic low speed | 18 |
| roads: high traffic high speed | 36 |
| roads: highways | 24 |
| roads: unclassified | 12 |
| linear features in construction | 6 |
| public transportation: trams | 6 |
| public transportation: railways | 12 |
| fences | 1 |
| riverways | 6 |

Table S4. Cities included in the validation analysis, the OSM ID of the polygons included and the buffer size used for generating the study area, which was conformed of an enveloped surrounding the OSM polygon of the city's boundary.

| | City | State | OSM ID | buffer |
|---|---|---|---|---|
| 1 | Wilmington | Delaware | 369472 | 15 |
| 2 | Edmonton | Alberta | 2564500 | 10 |
| 3 | Phoenix | Arizona | 111257 | 45 |
| 4 | Little Rock | Arkansas | 111147 | 10 |
| 5 | Vancouver | BC | 1852574 | 10 |
| 6 | Berkeley | California | 2833528 | 10 |

| | | | | |
|---|---|---|---|---|
| 7 | Pasadena | California | 3529725 | 10 |
| 8 | Pomona | California | 1532363 | 10 |
| 9 | Fort Collins | Colorado | 112524 | 20 |
| 10 | Atlanta | Georgia | 119557 | 35 |
| 11 | Chicago | Illinois | 122604 | 46 |
| 12 | Urbana | Illinois | 126133 | 25 |
| 13 | National Capital Area | Maryland | 162112 | 10 |
| 14 | Boston | Massachusetts | 2315704 | 35 |
| 15 | St Louis | Missouri | 1180533 | 30 |
| 16 | Manchester | New Hampshire | 305467 | 25 |
| 17 | New York | New York | 175905 | 35 |
| 18 | Syracuse | New York | 174916 | 15 |
| 19 | Albany | New York | 175549 | 15 |
| 20 | Raleigh | North Carolina | 179052 | 2 |
| 21 | Toronto | Ontario | 324211 | 10 |
| 22 | Peterborough | Ontario | 7473849 | 2 |
| 23 | Saskatoon | Saskatchewan | 4189345 | 10 |
| 24 | DFW | Texas | 115274 | 10 |
| 25 | Houston | Texas | 2688911 | 15 |
| 26 | Salt Lake City | Utah | 198770 | 35 |
| 27 | Woodbury_NY | New York | 175473 | 25 |
| 28 | San Diego | California | 396482 | 60 |
| 29 | San Jose CA | California | 112143 | 5 |
| 30 | Calgary | Alberta | 3227127 | 1 |
| 31 | Athens | Georgia | 119353 | 5 |
| 32 | Victoria | British Columbia | 2221062 | 10 |
| 33 | Key Largo | Florida | 11109 | 30 |

Table S5. Reclassification table between global landcover layer and the classes included in this study.

| CEC landcover class description | CEC | OSM |
|---|---|---|
| Temperate or sub-polar needleleaf forest | 1 | 11 |
| Sub-polar taiga needleleaf forest | 2 | 11 |
| Tropical or sub-tropical broadleaf evergreen forest | 3 | 11 |
| Tropical or sub-tropical broadleaf deciduous forest | 4 | 11 |
| Temperate or sub-polar broadleaf deciduous forest | 5 | 11 |
| Mixed forest | 6 | 11 |

| | | |
|---|---|---|
| Tropical or sub-tropical shrubland | 7 | 9 |
| Temperate or sub-polar shrubland | 8 | 9 |
| Tropical or sub-tropical grassland | 9 | 9 |
| Temperate or sub-polar grassland | 10 | 9 |
| Sub-polar or polar shrubland-lichen-moss | 11 | 9 |
| Sub-polar or polar grassland-lichen-moss | 12 | 6 |
| Sub-polar or polar barren-lichen-moss | 13 | 10 |
| Wetland | 14 | 9 |
| Cropland | 15 | 8 |
| Barren lands | 16 | 10 |
| Urban | 17 | 28 |
| Water | 18 | 12 |
| Snow and Ice | 19 | 12 |

Table S6. Building footprint source metadata information for each city where layer was available.

| | city | State | Source | Attribution | year |
|---|---|---|---|---|---|
| 1 | Chicago | Illinois | aerial photography | city of Chicago | 2015 |
| 2 | Fort Worth | Texas | aerial photography | city of Fort Worth | 2018 |
| 3 | New York | New York | aerial photography | city of New York | 2016 |
| 4 | Pomona | California | aerial photography | city of Los Angeles | 2014 |
| 5 | Raleigh | North Carolina | aerial photography | city of Raleigh | 2018 |
| 6 | San Jose | California | aerial photography | city of Cupertino | 2011 |
| 7 | Victoria | British Columbia | aerial photography | city of Victoria | 2019 |
| 8 | Urbana | Illinois | computer-generated from pixel data | city of Champaign | 2018 |
| 9 | Albany | New York | computer-generated from pixel data | Cornell University, Microsoft | 2018 |
| 10 | Salt Lake City | Utah | computer-generated from pixel data | Utah State, Microsoft | 2018 |
| 11 | Syracuse | New York | computer-generated from pixel data | Cornell University, Microsoft | 2018 |
| 12 | Toronto | Ontario | derived from landcover raster | City of Toronto | 2014 |
| 13 | Phoenix | Arizona | LiDAR | USGS | 2021 |

| | | | | | |
|---|---|---|---|---|---|
| 14 | St Louis | Missouri | ortho photography | St. Charles County | 2012 |
| 15 | Vancouver | British Columbia | ortho photography | city of Vancouver | 2015 |
| 16 | Boston | Massachusetts | ortho photography and LiDAR | Massachusetts government | 2021 |
| 17 | Pasadena | California | stereo imagery | city of Pasadena | 2019 |
| 18 | Edmonton | Alberta | aerial photography | city of Edmonton | 2019 |
| 19 | Woodbury_NY | New York | computer-generated from pixel data | Cornell University, Microsoft | 2018 |
| 20 | National Capital Area | Maryland | aerial photography | Washington DC | 2018 |

Table S7. Precision values across landcover classes in our framework's output integrating OSM features to a global landcover map

| | |
|---|---|
| Overall accuracy | 0.89 |
| Kappa statistic | 0.89 |
| Lower accuracy | 0.88 |
| Upper accuracy | 0.90 |
| P-value | < 2.2e-16 |

| | Precision | Recall | F1 | Balanced accuracy |
|---|---|---|---|---|
| Class: 1 | 0.94 | 0.96 | 0.95 | 0.98 |
| Class: 2 | 0.95 | 0.99 | 0.97 | 0.99 |
| Class: 3 | 0.98 | 0.99 | 0.98 | 0.99 |
| Class: 4 | 0.95 | 0.97 | 0.96 | 0.99 |
| Class: 5 | 0.90 | 0.95 | 0.93 | 0.98 |
| Class: 6 | 0.90 | 0.57 | 0.70 | 0.78 |
| Class: 7 | 0.95 | 1.00 | 0.97 | 1.00 |
| Class: 8 | 0.69 | 0.93 | 0.79 | 0.96 |
| Class: 9 | 0.84 | 0.67 | 0.74 | 0.83 |
| Class: 10 | 0.94 | 0.61 | 0.74 | 0.80 |
| Class: 11 | 0.94 | 0.76 | 0.84 | 0.88 |
| Class: 12 | 0.78 | 0.94 | 0.85 | 0.97 |
| Class: 13 | 1.00 | 0.82 | 0.90 | 0.91 |
| Class: 14 | 1.00 | 0.88 | 0.94 | 0.94 |
| Class: 15 | 0.96 | 0.99 | 0.97 | 0.99 |
| Class: 16 | 0.95 | 0.99 | 0.97 | 0.99 |

| | | | | |
|---|---|---|---|---|
| Class: 17 | 0.95 | 1.00 | 0.97 | 1.00 |
| Class: 18 | 0.95 | 0.99 | 0.97 | 0.99 |
| Class: 19 | 0.95 | 1.00 | 0.98 | 1.00 |
| Class: 20 | 0.97 | 0.83 | 0.89 | 0.91 |
| Class: 21 | 0.95 | 1.00 | 0.98 | 1.00 |
| Class: 22 | 1.00 | 0.98 | 0.99 | 0.99 |
| Class: 23 | 0.97 | 1.00 | 0.98 | 1.00 |
| Class: 24 | 0.94 | 0.90 | 0.92 | 0.95 |
| Class: 25 | 0.98 | 0.99 | 0.98 | 0.99 |
| Class: 26 | 0.56 | 1.00 | 0.72 | 0.99 |
| Class: 27 | 0.80 | 1.00 | 0.89 | 1.00 |
| Class: 28 | 0.35 | 1.00 | 0.52 | 0.99 |

### Supplementary Figures

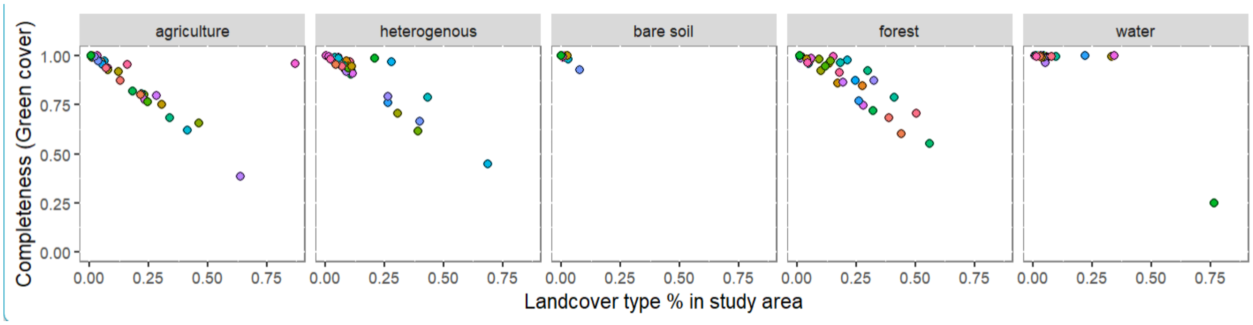


Figure S1. Completeness across green cover types with increasing surface area proportion of each landcover type

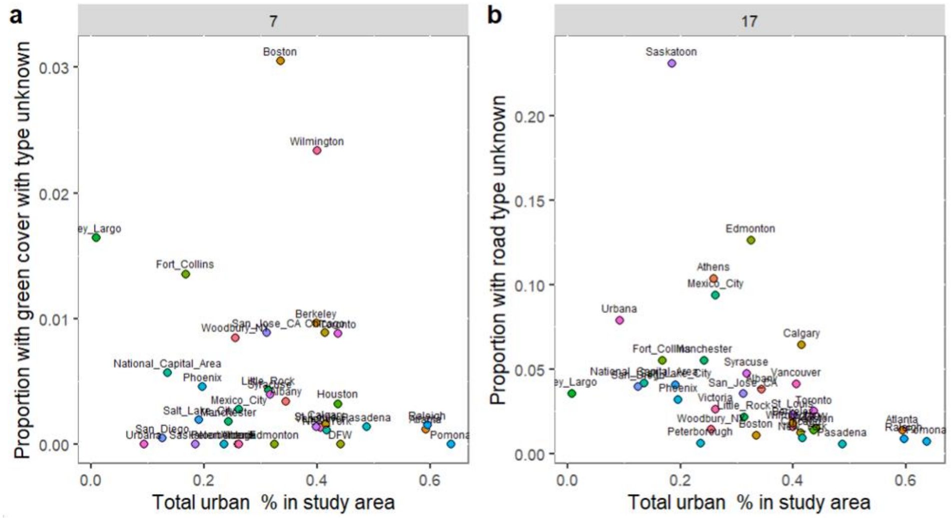


Figure S2. The proportion of landcover types including classes with unknown sub-typology, protected areas, class 7 (a) and roads, class 17 (b).

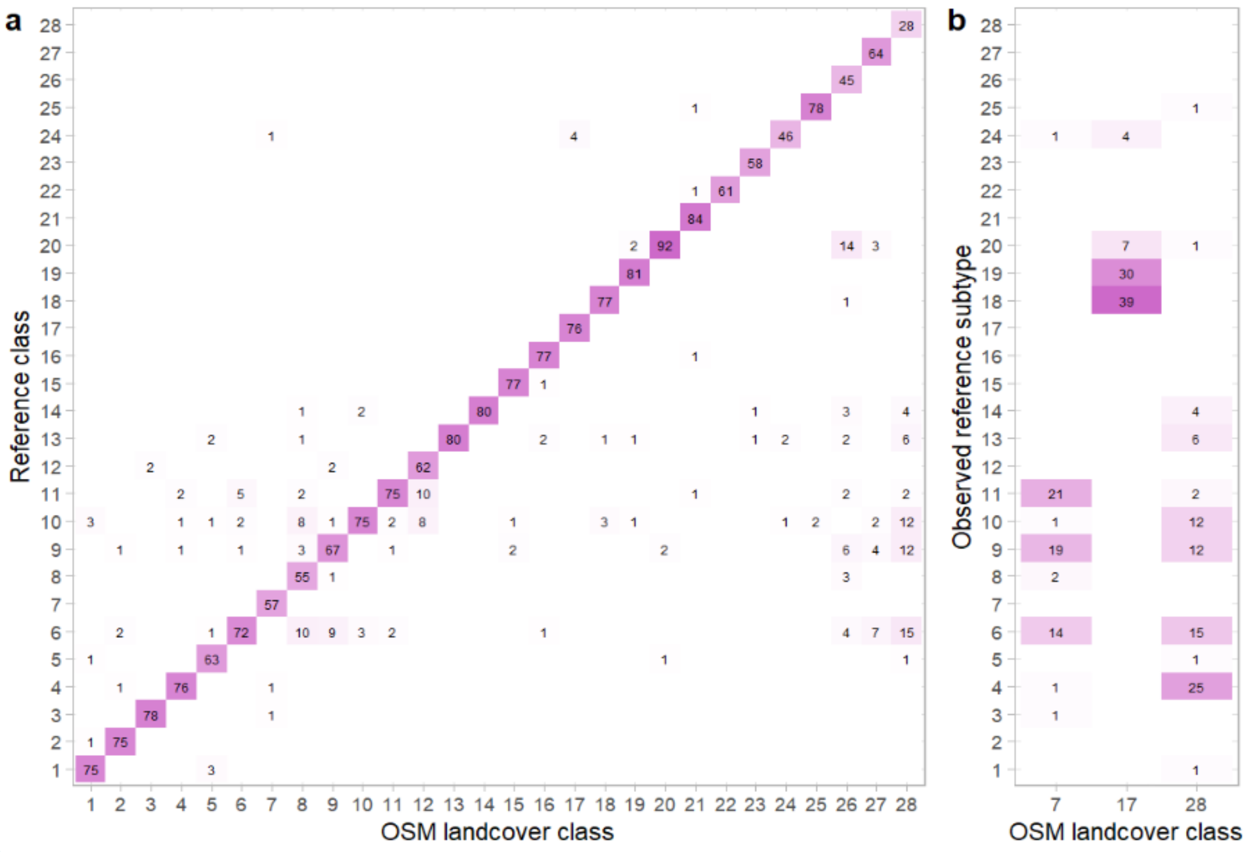


Figure S3. Confusion matrix showing the correspondence, across balanced random points, between the information provided by our framwork's output integrating OSM features into a continental scale landcover map and aerial photography and street view imagery (Google, 2023).

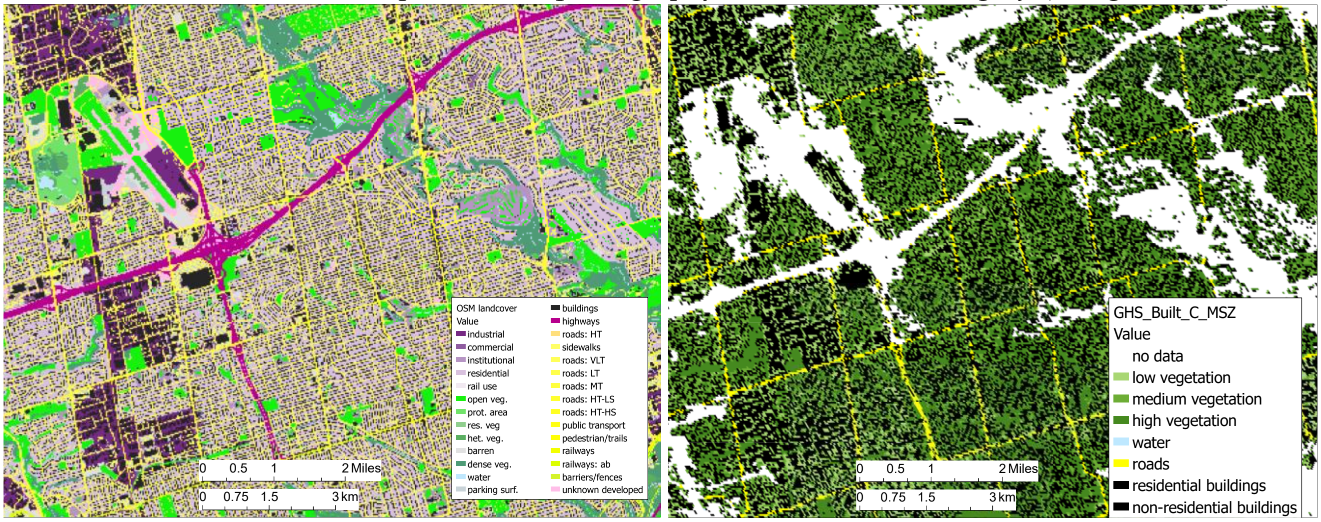


Figure S4. Comparison between our framework's output integrating OSM features and the Global human settlement (GHS) MSZ layer derived from remote sensing, resampled to 30m for comparison purposes.

Glossary

1. Land use: “term used to describe the human use of land. It represents the economic and cultural activities (e.g., agricultural, residential, industrial, mining, and recreational uses) that are practiced at a given place. Land use differs from land cover in that some uses are not always physically obvious.
2. Infrastructure: set of facilities and systems that serve a city and encompasses the services and facilities necessary for its economy, households, and firms to function. Infrastructure is composed of public and private physical structures such as roads, railways, bridges, tunnels, water supply,
3. Green cover: Green cover can be defined by all visible vegetation, natural or planted.
4. Land cover: the surface cover on the ground, whether vegetation, urban infrastructure, water, bare soil or other.
